# How does grasshopper *Oxya chinensis* respond to the artificially infected bacteria?

**DOI:** 10.1101/683680

**Authors:** Xiaomin Zhang, Keshi Zhang

## Abstract

*Oxya chinensis* is one of the most widespread grasshopper species found in China and one of the most common pests against rice. Due to the importance of haemocytes in insect immunity and limited information on the haemocytes of *O. chinensis*, their haemocytes were examined in detail. The cellular response of the grasshopper was challenged with bacteria *Escherichia coli*, *Staphylococcus aureus* and *Bacillus subtilis*. The morphology of the haemocytes was illustrated by the use of light, scanning electron and transmission electron microscopy, where different morphological varieties of haemocytes were observed. Granulocytes and plasmatocytes responded to the challenged bacteria by phagocytosis. The histochemical staining has indicated the presence of acid phosphatase in plasmatocytes and granulocytes. Non-phagocytic prohemocytes and vermicytes were noticed, but their functions in the circulation are unclear. Our results demonstrate an essential role of plasmatocytes and granulocytes in the innate immunity of *O. chinensis*. Insect haemocytes play a crucial role in cellular immunity, and further research is needed for a comprehensive understanding.

**Summary statement:** The cellular response was used by *Oxya chinensis* against the bacterial challenges where two types of haemocytes shared the duty of phagocytosis.

## 1. Introduction

Insect haemocytes have had scientists’ attention for many years since they were first discovered by Swammerdam (1637-1680) (Jones, 1962). Such attraction is because of the crucial role haemocytes play in insect immunity. Insects respond to pathogenic challenges through the interplay of two mechanisms, humoral and cellular defences (Charles and Killian, 2015, Urbański et al., 2018). Haemocytes mediate cellular defence by carrying out roles such as phagocytosis, nodulation, and encapsulation, but also facilitate humoral defence by synthesising and releasing enzymes, and other immune factors (Charles and Killian, 2015, Hillyer, 2016, Urbański et al., 2018).

Insect haemocytes are comprised of different populations which vary in their morphology and function (Strand, 2008). The origin, function, and classification of these haemocytes are debatable and not fully understood (Duressa and Huybrechts, 2016). Lack of a standardised protocol for insect immune investigation is one of the causes. Direct comparison of the insect immune response is difficult due to the variety of challenges used, life stages examined and methodologies practiced (Charles and Killian, 2015). The nomenclature of the haemocytes is not normalised across Insecta (Hillyer, 2016), but recent studies have attempted to generalise such variation. The most often reported insect haemocytes are prohemocytes, hypothetical small and spherical haemocyte progenitors, with a high nuclear to cytoplasm ratio; plasmatocytes, spindle-shaped cells with the presence of cytoplasmic projections and absence of granules in the cytoplasm, which function in phagocytosis and encapsulation; granulocytes (or granular cells), spherical and oval cells containing uniformly sized and electron-dense cytoplasmic granules, which acts as phagocytes; oenocytoids, large, uniformly shaped cells with a low nuclear to cytoplasm ratio and eccentrically located nucleus, which contain phenoloxidase and responsible for the haemolymph darkening (melanisation); and spherulocytes, irregular shaped cells packed with large inclusions (spherules), which may be a source of cuticular components (Ribeiro and Brehélin, 2006, Castillo et al., 2006, King and Hillyer, 2013, Grigorian and Hartenstein, 2013, Hillyer and Strand, 2014). Other occasionally found insect haemocytes include cystocytes (or coagulocytes), oval or fusiform cells usually lysed or degranulated in vitro, which differ from granulocytes under the periodic acid-Schiff reaction; adipohemocytes, spherical or oval cells, with the presence of reflective fat droplets and non-lipid granules and vacuoles; vermicytes (podocytes or vermiform cells), irregular shaped cells with multiple cytoplasmic extensions and small electron-dense granules; and megakaryocytes, large cells filled by the nucleus and minimum cytoplasm (Jones, 1977, Akai and Sato, 1979, Gupta, 1979, Gillespie et al., 2000, Ribeiro and Brehélin, 2006, Brehélin et al., 1978, Ren et al., 2014).

Grasshoppers play an essential role in the grassland ecosystem and have considerable economic impacts on agriculture (Latchininsky, 2013, Duressa et al., 2015). The grasshopper, *Oxya chinensis*, is the most widely distributed grasshopper species in China, and one of the major pests of rice, maize, sorghum and wheat (Liu et al., 1999, Zhang and Huang, 2008). Much attention of insect haemocytes has been paid to *Drosophila melanogaster* and selected lepidopteran species (Lavine and Strand, 2002). Hitherto, research on the *O. chinensis* haemocyte morphology was only from our early study, Wang et al. (2011), where three stains were employed to compare the staining efficiency. Thus, we aimed to provide a more comprehensive understanding of the grasshopper *O. chinensis* haemocytes and insights into standardising the insect haemocyte examination methods. The cellular response of *O. chinensis* was elicited by injection with live *Escherichia coli*, *Bacillus subtilis* and *Staphylococcus aureus*. The bacteria are frequently used in insect immune studies due to their non-pathogenicity for insects (Arp et al., 2017). Light microscopy was used with multiple stains to examine the haemocyte morphological and histochemical characteristics, and cellular response to bacterial challenges. Scanning and transmission electron microscopy were used to provide extra details of the haemocytes.

## 2. Materials and Methods

### 2.1 Species

Grasshopper *O. chinensis* adults were collected from Jinyuan District, Taiyuan, Shanxi Province, China (latitude: 37°42’24.51 "; longitude: 112°26’33.05 "; altitude: 779.72 meters). The collected samples were taken to the laboratory alive, kept in a well-ventilated room and fed with rice. They were maintained for a week before their blood sample was taken and treatment was done.

### 2.2 Microscopy

Haemolymph was extracted from the prothoracic leg bases of each grasshopper. The ethanol-sterilised leg base skin was pierced, and blood was taken with a pipette (MicroPette Plus). The blood smears were prepared for light (Olympus BX-51) and scanning electron microscopy (Hitachi S-570). For light microscopy, smears were prepared conventionally (more details) and stained with Wright-Giemsa dye and four histochemical procedures: acid phosphatase (ACP) reaction, periodic acid-Schiff (PAS) reaction, oil Red O staining and chloroacetate esterase (CAE) reaction. All dyes were purchased from Baso Diagnostics Inc. (Beijing, China) and smears were stained according to the manufacturer’s protocol.

For scanning electron microscopy, smears were fixed in 3% glutaraldehyde, washed with 0.1M phosphate (pH 7.4) and post-fixed with 1% osmium tetroxide. Fixed smears were then dehydrated in a graded series of ethanol (20%, 40%, 60%, 75%, 90%, 95%, and 100%) and followed by a graded series of tertiary butanol. Dehydrated smears were freeze-dried with the FD-1A-50 freeze dryer (Beijing, China) and coated with gold using the SBC-12 ion sputter coater (Shanghai, China).

For transmission electron microscopy (JEM-1400), the extruded blood was fixed in tubes with 5% glutaraldehyde. The tubes were centrifuged at 3000rpm for 5min, and the supernatant was discarded. Haemocyte pellets were post-fixed with 1% osmium tetroxide, and dehydrated in a graded ethanol series and then acetone before being embedded in Epon812 resin. Sections were cut with an ultramicrotome (Leica Rm2255) and stained with uranyl acetate and followed by lead citrate.

### 2.3 Bacterial infection experiment

Bacteria *E. coli*, *B. subtilis* and *S. aureus* were purchased from China General Microbiological Culture Collection Centre (Beijing, China). Thirty adult grasshoppers were separated into three groups and were injected, respectively, with 10μl of *E. coli*, *B. subtilis* or *S. aureu*or (1×10^5^ ml^−1^) suspended in phosphate buffered saline (pH7.4). Bacteria were injected using a microsyringe between the second and third ethanol (75%) sterilised segments of the abdomen. All grasshoppers were alive at the end of the experiment. The thorax of the grasshoppers was disinfected with ethanol and pierced with a needle to extract haemolymph at 4, 8, 12, 24 and 48 hours after the treatment. A gentle squeeze of the thorax was applied to promote bleeding, and a pipette was used to collect blood. Slides were prepared and stained with Wright-Giemsa as described previously.

### 2.4 Haemocyte count and statistical methods

Different haemocyte types were counted from the Wright-Giemsa stained slides, and their relevant percentage was calculated. Ten non-treated grasshoppers were selected, and at least four hundred cells were counted from each. The number of phagocytic haemocytes of the bacterial treated grasshoppers was counted at 4, 8, 12, 24 and 48 hours after the injection. Statistica 10 data analysis software (StatSoft, 2011) was used to run statistical tests and generate graphs. Data obtained from the differential count was analysed through ANOVA and Tukey’s Honestly Significant Difference test (p<0.05).

## 3. Results

### 3.1 Haemocyte morphology

The haemocytes of *O. chinensis* varied regarding their shape, size and cytoplasmic contents. They responded to the bacterial challenges by phagocytosis (Fig 1). The phagocytes bound the bacteria *E.coli* and *B. subtilis* and *S. aureus*, and digested them internally. The phagocytes were capable of digesting the engulfed bacteria while attaching new ones. From four to eight hours after the injection, most bacteria were attached to the haemocyte membrane while some was being ingested. After12 hours, most bacteria inside the phagocytes were broken down into fractions. However, the binding of bacteria by phagocytes was observed at the end of the experiment. Thus, complete removal of the injected bacteria by *Oxya*’s immune system requires more than two days. The bacteria *B. subtilis* formed endospores as showing by their transparent centre were digested by the phagocytes of *O. chinensis*.

**Fig. 1.**
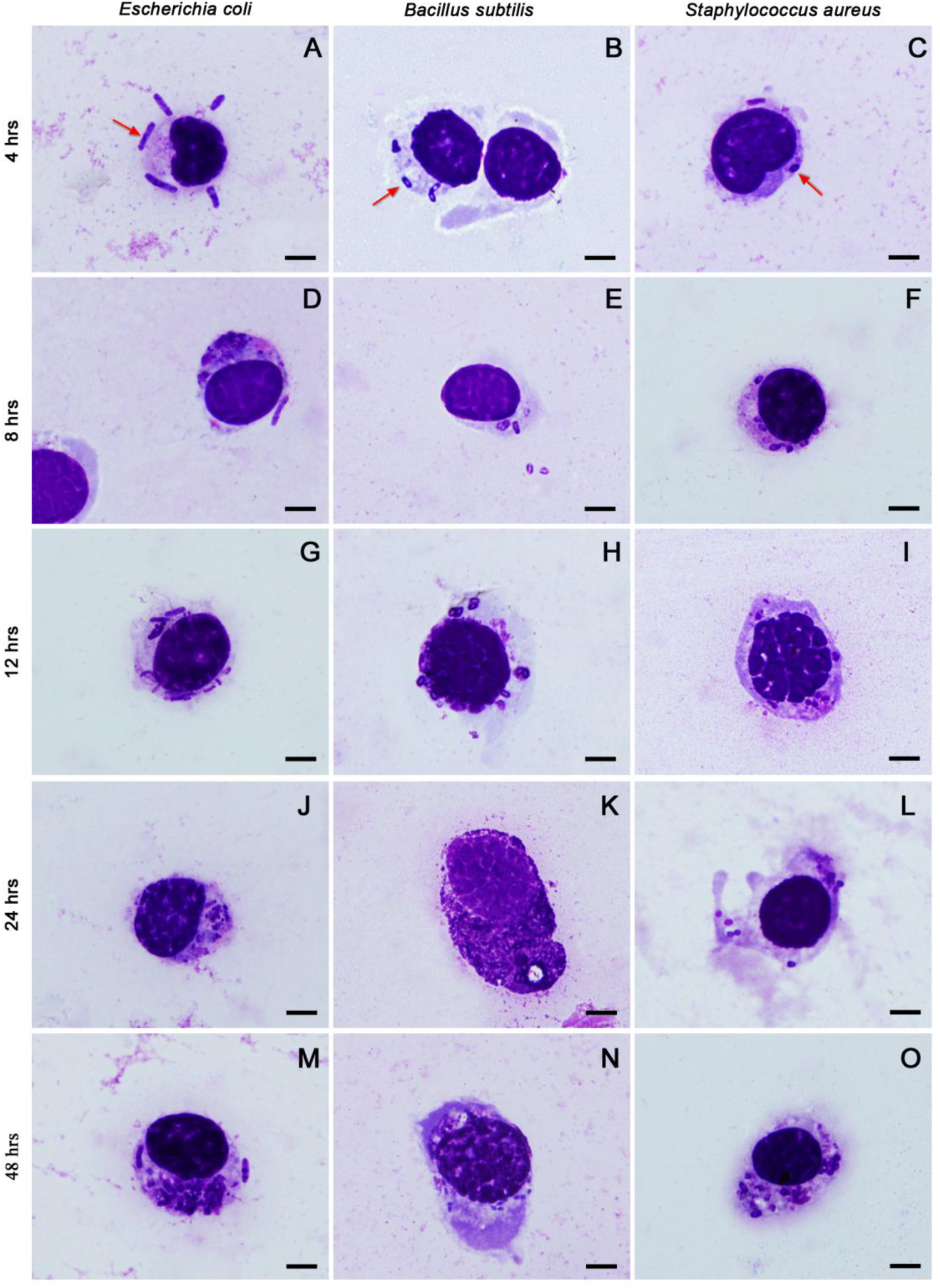
The phagocytic response of grasshopper *Oxya chinensis* against bacteria *Escherichia coli*, *Bacillus subtilis* and *Staphylococcus aureus* at 4, 8, 12, 24 and 48 hours post-injection; the phagocytes showing are plasmatocytes (A-J, L-O) and granulocyte (K); arrows point to bacteria; bars denote 5μm.

The phagocytes of *O. chinensis* contained two morphological varieties, which were distinguished by the existence of cytoplasmic granules. Round, oval and irregular shaped granulocytes contained small (<1µm) basophilic granules (purple) were measured 12-34µm in diameter (Fig. 2). The centrally located nucleus was stained dark red or bluish purple with Wright-Giemsa measured 7-20µm in diameter. The cytoplasm of the granulocytes was stained transparent (neutral), pink (eosinophilic) or light blue (basophilic). The density of the granules ranged from few to packed, and the distribution was uniform or erratic.

**Fig. 2.**
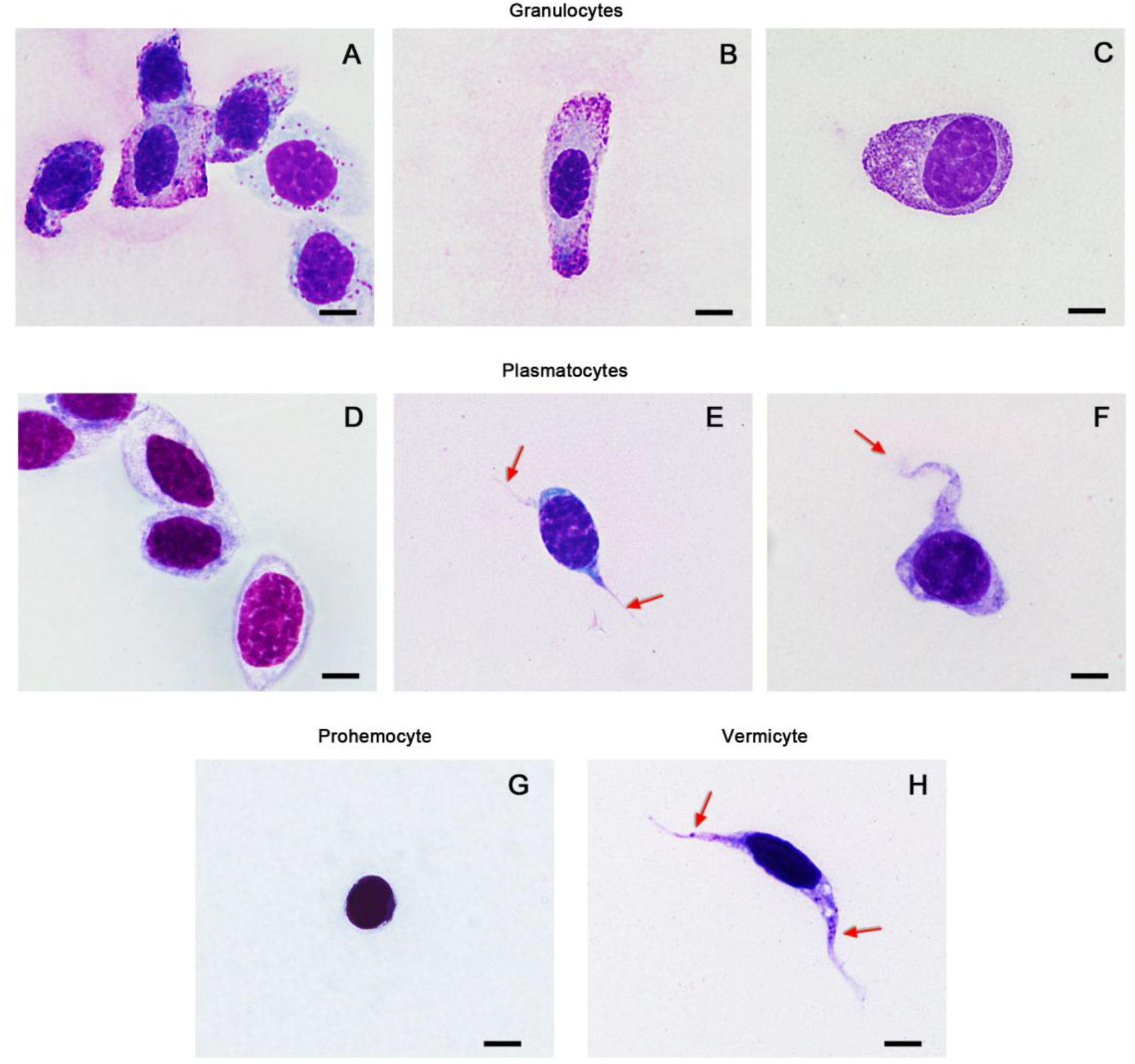
Wright-Giemsa stained haemocytes of *Oxya chinensis* under the light microscope: polymorphic granulocytes (A-C) with purple cytoplasmic granules which varies in density; oval-shaped plasmatocytes (A) and plasmatocytes (B & C) with extended pseudopodia (arrows); a spherical shaped prohemocyte with limited cytoplasm (F); a worm-shaped vermicyte (G) with elongated cytoplasm and nucleus contained some cytoplasmic granules (arrows); bars denote 5 μm.

Under the scanning electron microscope, granulocytes showed a rough surface (Fig. 3). The small lumps on the cell membrane were thought to be the cytoplasmic granules causing such embossment. These spherical swellings were used to distinguish granulocytes from the other haemocytes. Under the transmission electron microscope, the electron-dense and electron-lucent granules in the granulocyte were round or oval, and structure-less (Fig. 4). In addition to the granules, mitochondria, endoplasmic reticulum, phagosome-like vacuole and clumps of small bright inclusions were also found (Fig. 4D).

**Fig. 3.**
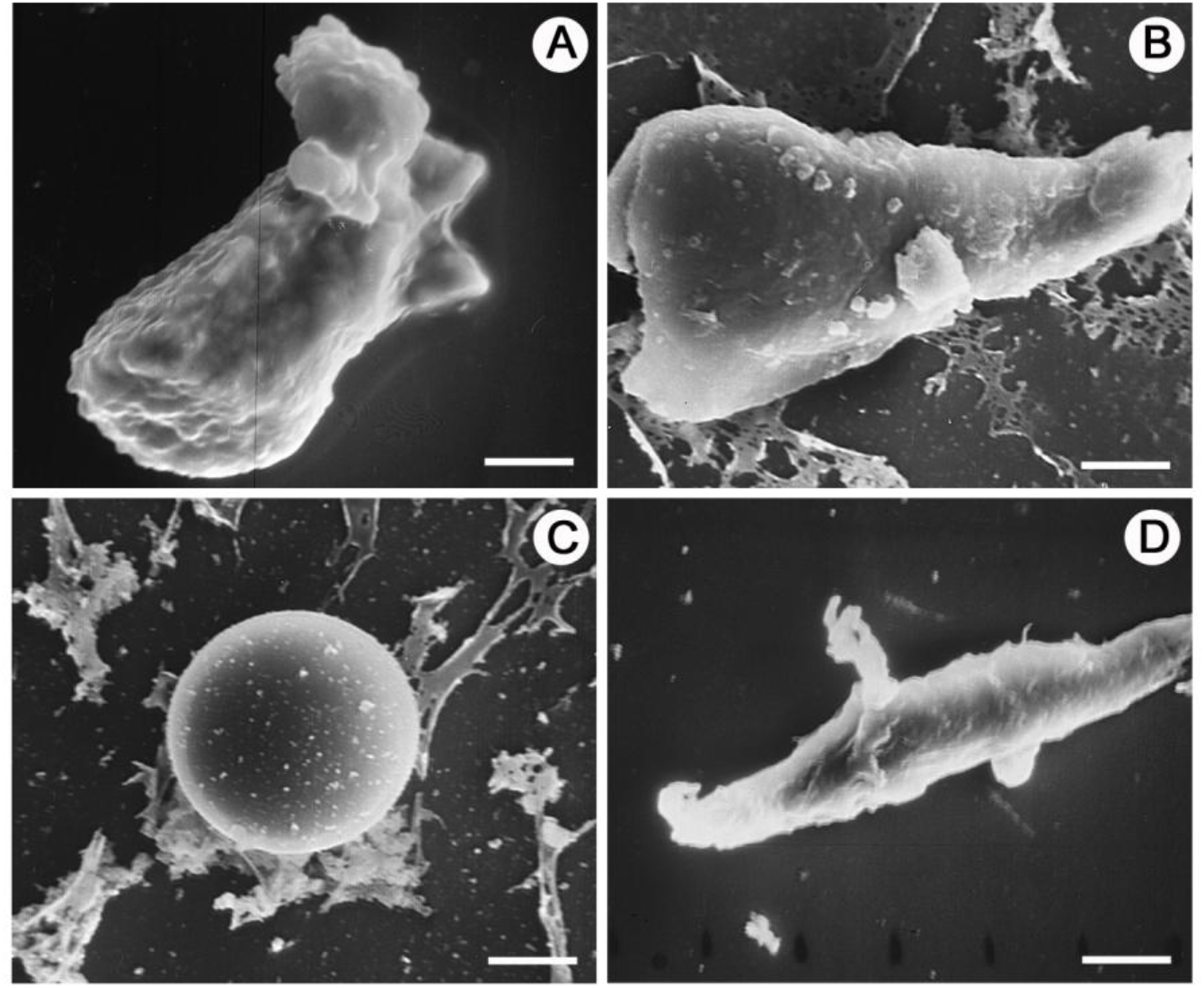
Haemocytes of *Oxya chinensis* under the scanning electron microscope: an irregular shaped granulocyte with lumps on the membrane caused the presence of cytoplasmic granules (A), bar denotes 7 μm; an irregular shaped plasmatocyte with a relatively smooth membrane (B), bar denotes 5 μm; a spherical shaped prohemocyte with smooth membrane (C), bar denotes 4 μm; a vermicyte with rough membrane and pseudopodia (D), bar denotes 7 μm.

**Fig. 4.**
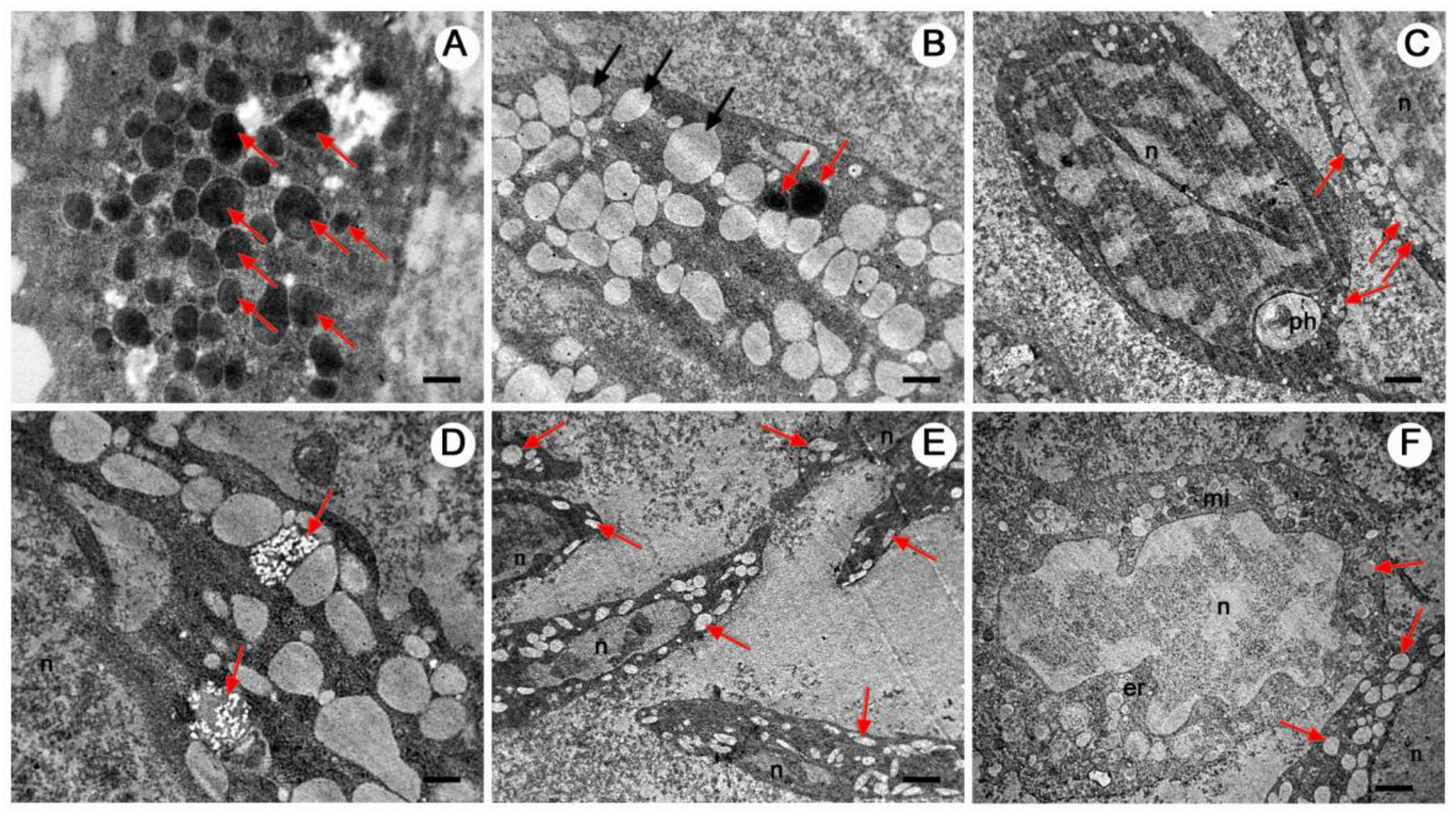
Granulocytes of *Oxya chinensis* under the transmission electron microscope: sections of granulocytes with visible electron-dense (red arrow) or electron-lucent (black arrow) granules in the cytoplasm (A & B); A: bar denotes 0.5 μm; B: bar denotes 0.8 μm; C: arrows points to some of the large and small electron-lucent granules in a granulocyte with a sizeable phagosome-like vacuole (ph), and section of a granulocyte, bar denotes 2 μm; D: section of a granulocyte with two clusters of small bright inclusions (arrows), bar denotes 0.6 μm; E: the elongated granulocytes are filled with many electron-lucent granules (arrows), bar denotes 2 μm; F: arrows indicate some of the electron-lucent granules, bar denotes 1 μm; nucleus (n), mitochondria (mi) and endoplasmic reticulum (er) are found in the cytoplasm.

The granular-less phagocyte, plasmatocytes, were polymorphic with round, oval, spindle and irregular shapes measured 10-32 μm in diameter (Fig. 2). Cytoplasmic projections including pseudopodia were seen under the light, scanning electron and transmission electron microscopes (Figs. 3 & 5). The nucleus of plasmatocytes was stained blue or purplish-red with Wright-Giemsa measured 8-18 μm in diameter. The cytoplasm of plasmatocytes was uniformly distributed around the nucleus and stained blue, pinkish red or colourless. Under the scanning electron microscope, the outer surface of plasmatocyte cell membranes was relatively smooth. Under the transmission electron microscope, mitochondria, endoplasmic reticulum, Golgi apparatus, vacuoles and other inclusions were recognised in the cytoplasm of the plasmatocyte (Fig. 5). Exocytosis-like activities were observed (Fig. 5B).

**Fig. 5.**
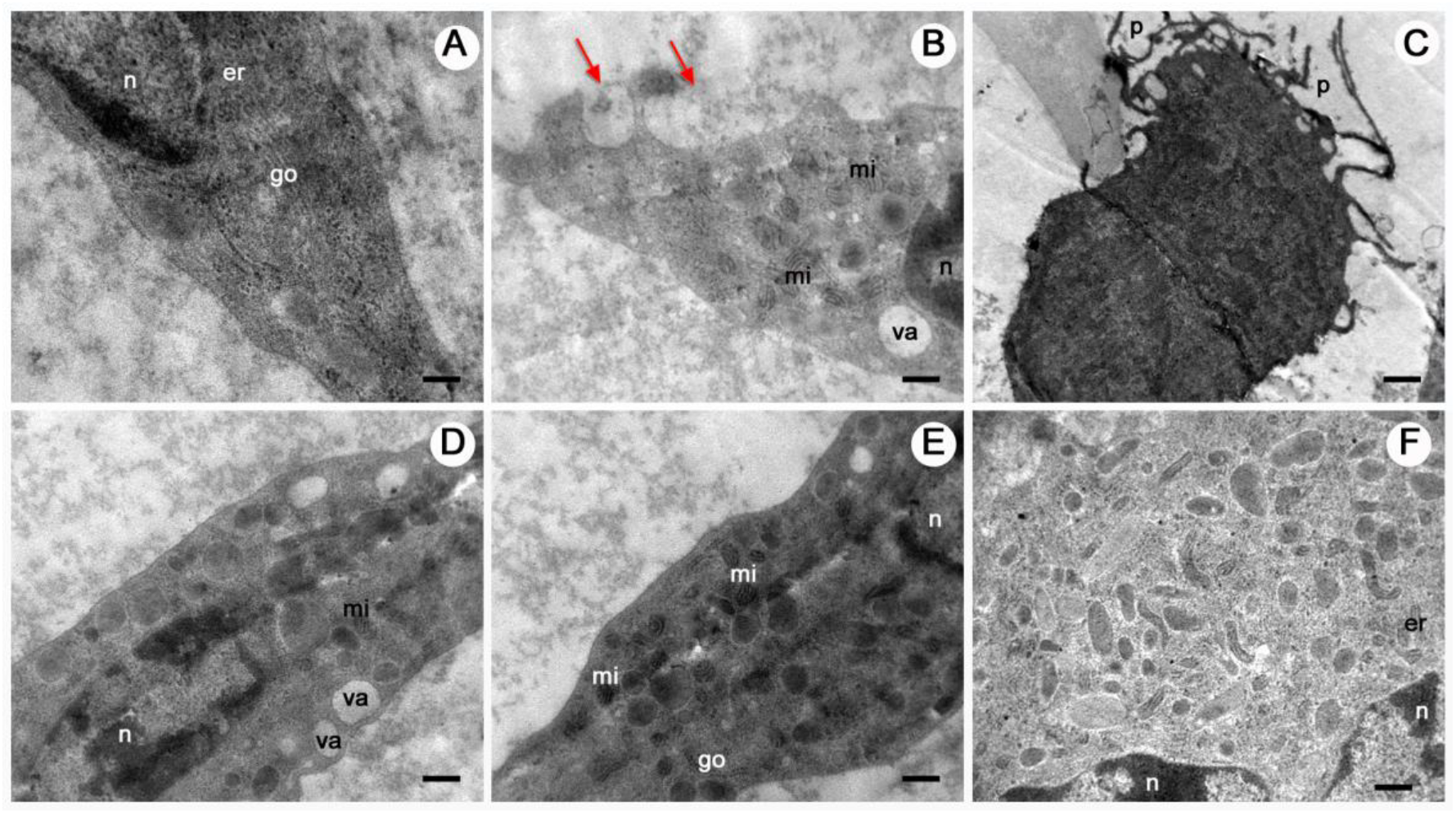
Plasmatocytes of *Oxya chinensis* under the transmission electron microscope: (A & B) bars denote 0.25 μm; (B) arrows indicate exocytosis-like activity; (C) many elongated pseudopodia (p) are extended outward of the plasmatocyte, bar denotes 1 μm; (D-F) the presence of nucleus (n), mitochondria (mi), endoplasmic reticulum (er), Golgi apparatus (go) and vacuole (va) is found in the cytoplasm; bars denote 0.5 μm.

Two non-phagocytic varieties of haemocytes were sporadically observed in this study. They were prohemocytes and vermicytes. These two cell types were morphologically distinctive to plasmatocytes and granulocytes. Prohemocytes were much smaller than the others, which displayed a spherical shape of approximately 8-12 µm in diameter (Fig. 2). The large nucleus was dyed purple or violet with Wright-Giemsa, which almost filled the whole cell. Prohemocytes had a smooth cell membrane under the scanning electron microscope with no pseudopodia or other cytoplasmic projections (Fig. 3). Under the transmission electron microscope, mitochondria, endoplasmic reticulum, Golgi apparatus, vacuoles, and other electron-dense and electron-lucent particles were observed in the cytoplasm of prohemocytes (Fig. 6).

**Fig. 6.**
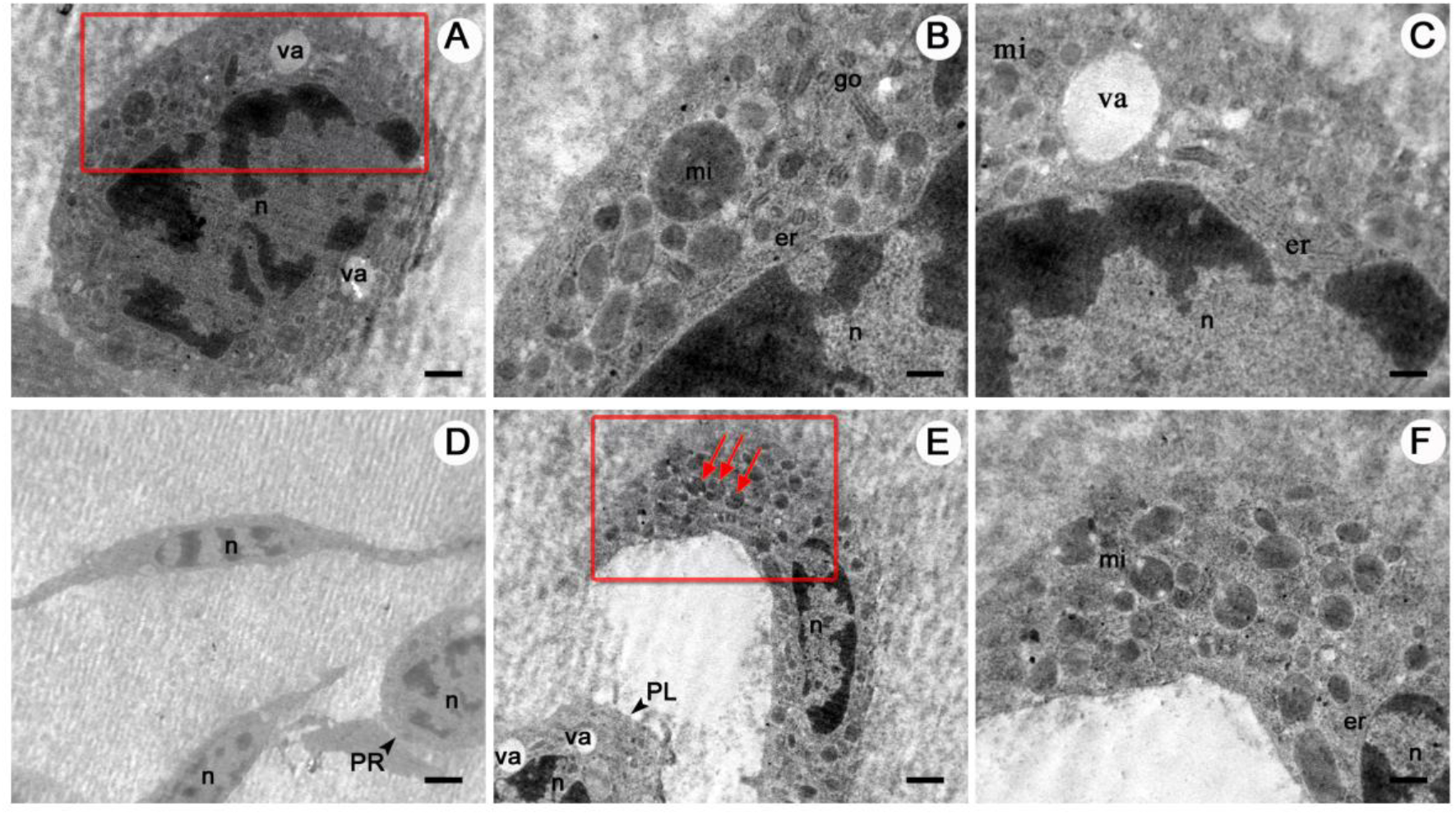
A prohemocyte of *Oxya chinensis* under the transmission electron microscope (A-C): B & C are enlarged sections of A (red rectangle); A: bar denotes 1 μm; B & C: bars denote 0.5 μm; nucleus (n), mitochondria (mi), endoplasmic reticulum (er) and vacuole (va) are found in the cytoplasm; vermicytes of *Oxya chinensis* under the transmission electron microscope (D-F); D) two worm-shaped vermicytes with elongated cytoplasm and nucleus, bar denotes 2.5 μm; E: arrows indicate some of the vermicytes’ cytoplasmic inclusions, bar denotes 1 μm; F: an enlarged section of E (red rectangle), bar denotes 0.5 μm; nucleus (n), mitochondria (mi), endoplasmic reticulum (er), vacuole (va), and additional prohemocyte (PR) and plasmatocyte (PL) were noticed.

Vermicytes or vermin-formed haemocytes were found with distinct worm-shaped cytoplasm and elongated nucleus (Fig. 2). The span of vermicyte extensions can be over 30 μm. Unlike plasmatocytes with elongated pseudopodia, the nucleus of vermicytes was attenuated and elongated with the cell. The dark blue nucleus stained by Wright-Giemsa was 8-25 μm in diameter. Granules were found in the cytoplasm of many vermicytes examined. Under the transmission electron microscope, the cytoplasm of vermicytes was abundant in the endoplasmic reticulum, mitochondria and other inclusions (Fig. 6). Vermicytes were not observed under the scanning electron microscope (Add vermicytes scanning photo).

### 3.2 Haemocyte histochemistry

Plasmatocytes and granulocytes were found to contain acid phosphatase (ACP) with a purplish-red colouration of the cytoplasm (Fig. 7). ACP-negative plasmatocytes and granulocytes were also observed with a clear cytoplasm or cytoplasm filled with clear refractive granules. Chloroacetate esterase (CAE) staining indicated the presence of positively reacted compound in the grasshopper’s haemomph and outside the haemocytes. No positively reacted lipid droplets were revealed by the oil red O staining (ORO) inside haemocytes. The ORO stained cytoplasm of some haemocytes was contrastively darker than others. Few haemocytes contained positively reacted pinkish-red substance with periodic acid-Schiff staining.

**Fig. 7.**
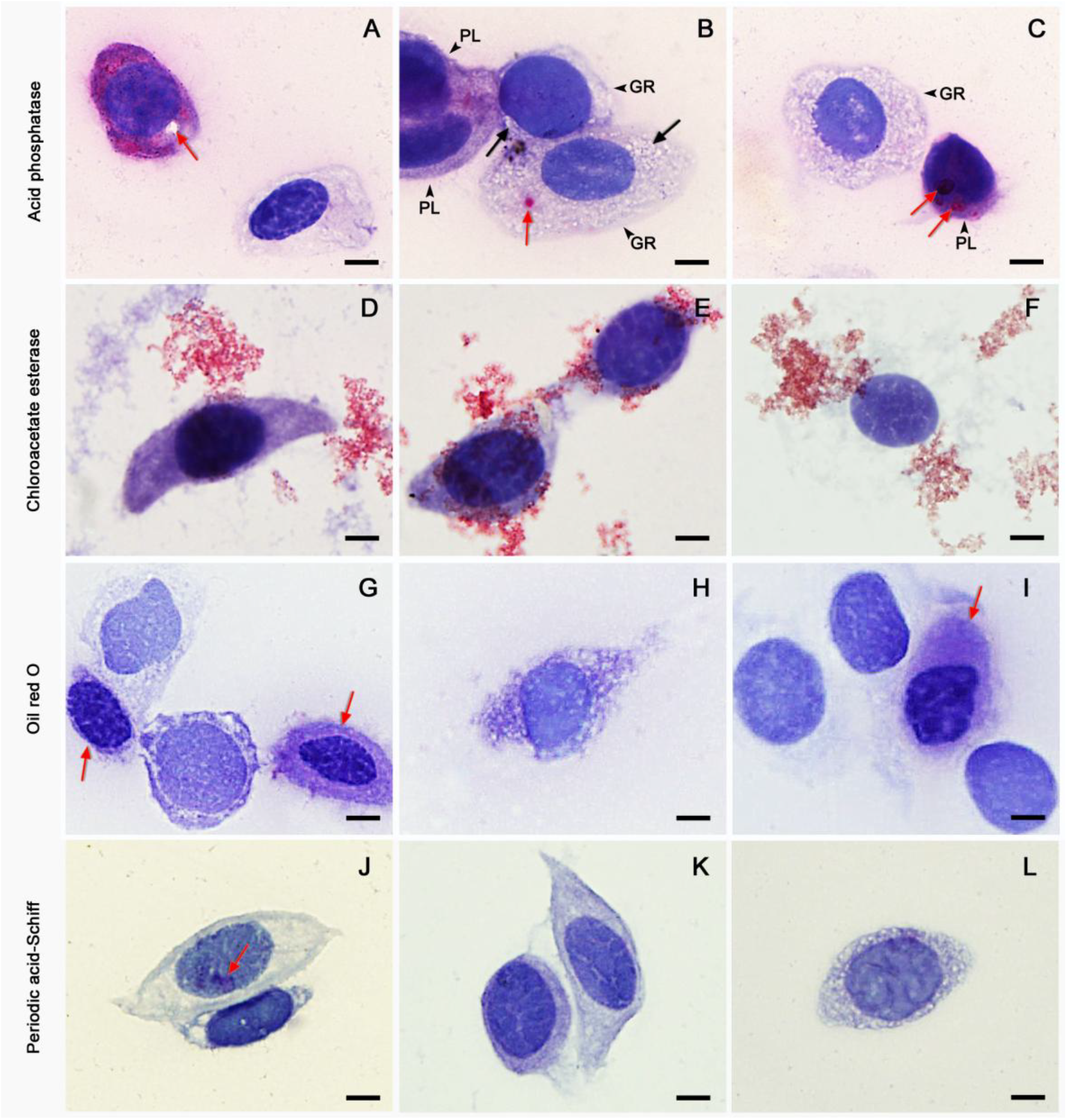
Histochemically stained haemocytes of *Oxya chineneis*: haemocytes stained with acid phosphatase reaction (ACP), positively reacted substance are purplish-red in colour (A-C); A: two plasmatocytes with one positively reacted to ACP reaction (purplish-red colouration) which contains an unstained vacuole (arrow); B: two plasmatocytes with weak positive reaction to ACP (black arrows) and two granulocytes with one contains an ACP-positive red granule (arrow); C: an ACP-positive plasmatocyte (arrow) and an ACP-negative granulocyte; haemocytes stained with chloroacetate esterase reaction, all positively reacted red substances are background staining (D-F); D & E: plasmatocytes; F: a naked nucleus with no conspicuous cytoplasm; haemocytes stained with Oil Red O staining (ORO; G-I); G) plasmatocytes with two weakly reacted to ORO (arrows); H: a negatively reacted granulocyte with clear granules; I: a weak ORO-positive plasmatocyte (arrow) and three nuclei with no defined cytoplasm; haemocytes stained with Periodic acid-Schiff (PAS) reaction (J-L); J: one of the two plasmatocytes has PAS-positive purplish-red substances (arrow); K: two negatively reacted plasmatocytes; L: a negatively reacted granulocyte with clear granules; bars denote 5 μm.

### 3.3 Differential haemocyte count

Plasmatocyte and granulocyte were the most common haemocytes found during the study, which comprised approximately 90 percent of the total cell counted (Fig. 8). Slightly more granulocytes were found than plasmatocytes in the untreated grasshopper samples. However, in the bacteria treated samples, significantly more plasmatocytes were found than the granulocytes especially with grasshoppers injected with *E. coli* and *S. aureus*. The number of plasmatocytes and granulocytes of the *B. subtilis* and *S. aureus* treated individuals varied during the 48-hour-period. The plasmatocytes and granulocytes changes were different across the three bacterial treated groups.

**Fig. 8.**
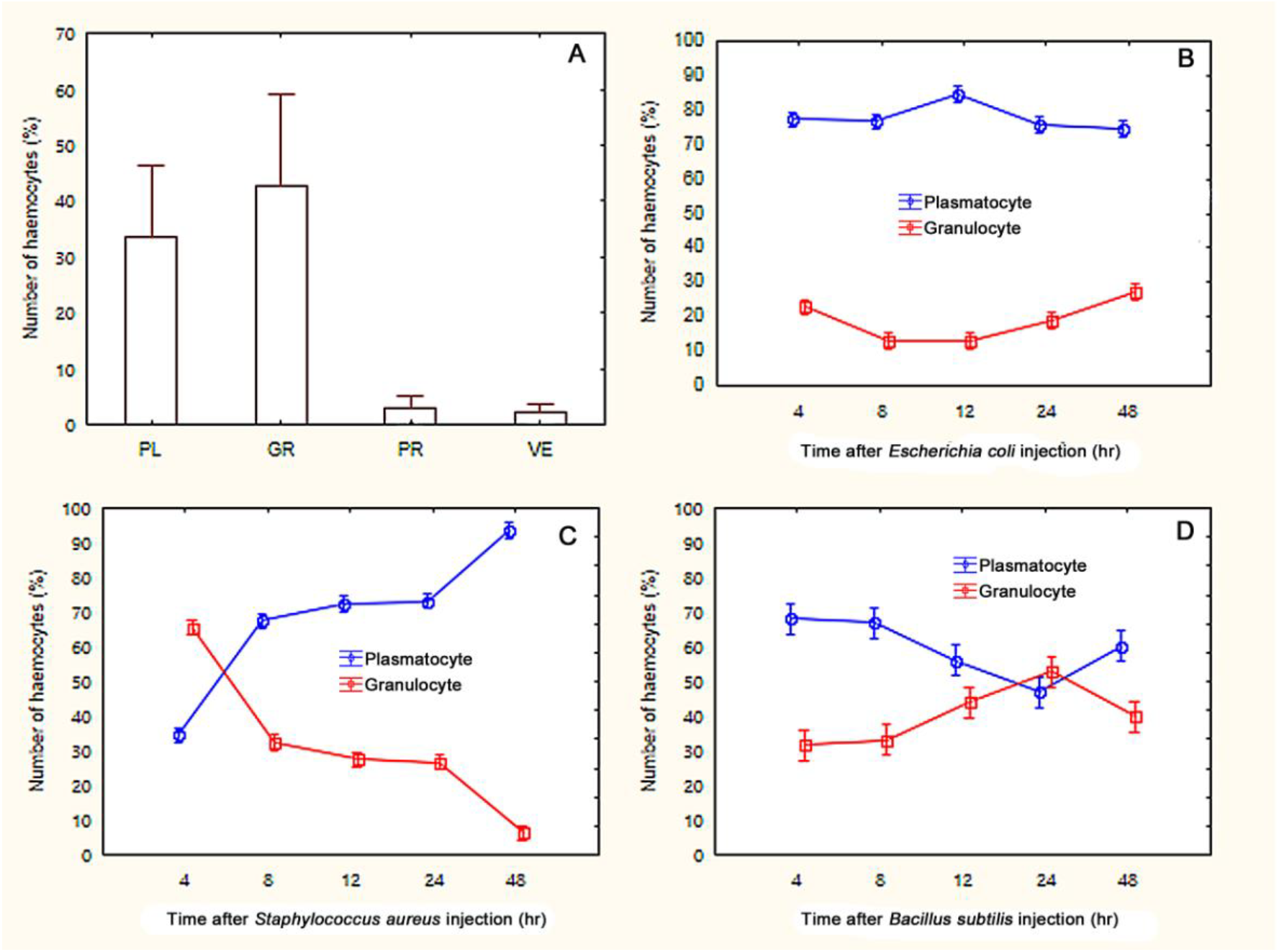
Haemocyte counts from the Wright-Giemsa stained preparations of normal adults (A) and adults injected with (B) *Bacillus subtilis*, (C) *Escherichia coli* and (D) *Staphylococcus aureus*. A: prohemocyte (PR), plasmatocyte (PL), granulocyte (GR) and vermicyte (VE); B-D: the number of plasmatocytes and granulocytes with and without the attachment or engulfment of the injected bacteria are included in the graph.

## 4. Discussion

Haemocytes of *O. chinensis* can be classified functionally as phagocytes and non-phagocytic haemocytes or morphologically as granulocytes, plasmatocytes prohemocytes and vermicytes. Phagocytes or more specifically granulocytes and plasmatocytes account for the majority of haemocytes observed in this study. Phagocytosis is the primary defensive strategy utilised by insects against bacterial infection, in which the phagocytes account for a substantial proportion of the insect haemocytes (Blandin and Levashina, 2007, Hillyer, 2016). The capacity in digesting endocytosed materials of *Oxya*’s plasmatocytes and granulocytes was indicated by the finding of the hydrolase, acid phosphatase. This hydrolase, is located in the lysosomes of haemocytes, and responsible for endogenous and exogenous macromolecules including the breaking down the bacteria cell wall (Callewaert and Michiels, 2010, Wu and Yi, 2015). Additionally, insect haemocytes release acid phosphatase into the haemolymph to modify the pathogens’ surface molecular structure thus to enhance the phagocytosis responses of haemocytes (Callewaert and Michiels, 2010).

The phagocytes of *O.chinensis* were morphological and functionally similar to most other insects. Both granulocytes and plasmatocytes were suggested as the first lines of the insect cellular defence which primarily involves in phagocytosis and encapsulation in many species (Grigorian and Hartenstein, 2013, Ribeiro and Brehélin, 2006, Hillyer, 2016, Tojo et al., 2000). The high abundance of plasmatocytes and granulocytes is observed in other grasshoppers and locusts (Anggraeni et al., 2011, Yu et al., 2016). Conversely, the structured granules of granulocytes observed under the TEM (Ribeiro and Brehélin, 2006) were not seen in this study. Granulocytes were mostly reported smaller than plasmatocytes being between 5-9 um (Ribeiro and Brehélin, 2006, Hillyer and Strand, 2014), but in *Oxya* were similarly sized to plasmatocytes. In Lepidoptera, plasmatocytes mostly exhibit in encapsulation rather than phagocytosis (Ribeiro and Brehélin, 2006).

Granulocytes of *Locusta migratoria* were found to contain lysozyme in the granules (Zachary and Hoffmann, 1984). However, in *Oxya*, the granules of granulocytes did not elicit the phagocytic ability of the haemocytes. Both plasmatocyte and granulocyte were found to participate in phagocytosis of the injected bacteria.

Our findings agree that not all plasmatocytes or granulocytes are involved in direct phagocytosis (Yu et al., 2016). Some plasmatocytes and granulocytes of *O. chinensis* contained no acid phosphatase and did not respond to the intruders. Granulocytes are reported to participate in the activation of the encapsulation response, which release opsonin-like materials in the presence of foreign objects (Browne et al., 2013, Hillyer, 2016). These materials cause the deformation of plasmatocytes to form multiple layers around the unwanted large substances into capsules (Ribeiro and Brehélin, 2006). Other functions of granulocytes such as the haemolymph clotting, wound healing and melanisation were reported in the greater wax moth *Galleria mellonella*, migratory locust *L. migratoria* and mosquitoes (Akai and Sato, 1979, Dushay, 2009, Hillyer and Strand, 2014).

Vermicytes were distinguished from plasmatocytes or granulocytes based on their morphology, histochemistry and function. The small inclusions of vermicytes and their non-phagocytic capacity have been used to separate them from the other haemocytes (Ribeiro and Brehélin, 2006). Most studies consider vermicytes as a sub-class or variant form of the plasmatocytes, which they are rarely reported in insects (Jones, 1977, Gupta, 1979, Ribeiro and Brehélin, 2006).

None of the *O. chinensis* haemocytes were morphologically corresponding to adipohemocyte, oenocytoid, podocyte and megakaryocyte. The oil red staining (ORO) found no lipid droplets in the haemocytes of *O. chinensis* thus an absence of adipohemocytes. ORO reveals neutral lipids such as triglycerides and cholesterol esters (Tame et al., 2018). Under the transmission electron microscope, the lipid droplets measuring about 1 μm in diameter filled the adipohemocytes of the Chinese grasshopper *Acrida cinerea* were not observed in *O. chinensis* (Yu et al., 1977).

In this study, cystocytes or coagulocytes were not considered as a distinct haemocyte type of *O. chinensis*. Our results are in contrast to those of Costin (1975) in separating cystocytes from plasmatocytes and granulocytes histochemically with the periodic acid-Schiff stain. The structured granules or globules of *Locusta migratoria* found under the transmission electron microscope were not seen in *O. chinensis* (Brehélin et al., 1978). Early studies have described cystocytes or coagulocytes as having fewer granules than granulocytes, and interpret the cytoplasmic remnants as coagulum or coagulated haemolymph (Costin, 1975, Gupta, 1979, Gillespie et al., 2000), while later studies have not mentioned their existence as a separate type. Cystocytes of *Locusta migratoria* is classified as a subpopulation of granulocytes, which contain less red granules and participated in phagocytosis (Yu et al., 2016). The most reported role of cystocytes is the coagulation of the insect haemolymph (Gupta, 1979, Zachary and Hoffmann, 1984). However, the coagulation process has been reported to require the cooperation of haemocytes and the plasma (Dushay, 2009). Granulocytes in many species have been suggested to function in coagulation (Grigorian and Hartenstein, 2013, Hillyer and Strand, 2014).

More plasmatocytes of *O. chinensis* were found than granulocytes in the bacteria infected samples, which differs from results with *O. japonica* where more granulocytes were found (Anggraeni et al., 2011). However, these may all be due to the difference in challenges used between the two studies. During the cellular response, plasmatocytes and granulocytes of *O. chinensis* were found to replenish or enhance their abundance. The increase in granulocytes and plasmatocytes of *Locusta migratoria* after inoculation of the fungus *Metarhizium acridum* was thought to result from a release of sessile haemocytes into the circulation (Yu et al., 2016). However, mitosis of circulating haemocytes is suggested as the primary source for haemocyte replenishment during the infection of pathogens in both mosquitoes and locusts (King and Hillyer, 2013, Duressa et al., 2015).

Based on our observations on *O. chinensis* and from other reported studies, the morphology and histochemistry of the haemocytes are essential in understanding their function. Some haemocytes could solely be an intermediate stage of the others. Plasmatocytes and granulocytes are the primary phagocytic cells of *O. chinensis* based on their abundance, morphology, histochemistry and phagocytic responses. The histochemical stains indicated that the haemocytes of *O. chinensis* are capable of digesting foreign materials, detoxicating the haemolymph and regulating the metabolism. The importance of the non-phagocytic haemocytes found in this study to the cellular response of *O. chinensis* is questionable due to their low abundance and unknown histochemical properties. Based on our results, we believe the light microscopic observation with stained preparations is still convincing in examining the morphology, histology, and function of insect haemocytes. Results of this study may contribute to the improvement and standardisation of the staining techniques used in the insect haemocyte studies. Morphological features alone are not sufficient to confirm the classification of insect haemocytes. Combinations of techniques are needed to confirm the haemocyte types and function.

## Acknowledgement

We are grateful to the anonymous reviewers for their valuable comments. This work was supported by the National Natural Science Foundation of China [grant number 30770239]; and the Natural Science Foundation of Shanxi Province China [grant number 2009011048].

## Reference

Akai, H. & Sato, S. 1979. Surface and internal ultrastructure of hemocytes of some insects. In: Gupta, A. P. (ed.) Insect Hemocytes, development, forms, functions and techniques. Cambridge, London: Cambridge University Press.

Anggraeni, T., Melanie & Putra, R. E. 2011. Cellular and humoral immune defenses of Oxya japonica (Orthoptera: Acrididae) to entomopathogenic fungi Metarhizium anisopliae. Entomological Research, 41, 1–6.

Arp, A. P., Martini, X. & Pelz-Stelinski, K. S. 2017. Innate immune system capabilities of the Asian citrus psyllid, Diaphorina citri. Journal of invertebrate pathology, 148, 94–101.

Blandin, S. A. & Levashina, E. A. 2007. Phagocytosis in mosquito immune responses. Immunological reviews, 219, 8–16.

Breh Lin, M., Zachary, D. & Hoffmann, J. A. 1978. A comparative ultrastructural study of blood cells from nine insect orders. Cell and tissue research, 195, 45–57.

Browne, N., Heelan, M. & Kavanagh, K. 2013. An analysis of the structural and functional similarities of insect hemocytes and mammalian phagocytes. Virulence, 4, 597–603.

Callewaert, L. & Michiels, C. W. 2010. Lysozymes in the animal kingdom. Journal of biosciences, 35, 127–160.

Castillo, J., Robertson, A. & Strand, M. 2006. Characterization of hemocytes from the mosquitoes Anopheles gambiae and Aedes aegypti. Insect biochemistry and molecular biology, 36, 891–903.

Charles, H. M. & Killian, K. A. 2015. Response of the insect immune system to three different immune challenges. Journal of insect physiology, 81, 97–108.

Costin, N. M. 1975. Histochemical observations of the haemocytes ofLocusta migratoria. The Histochemical Journal, 7, 21–43.

Duressa, T. F. & Huybrechts, R. 2016. Development of primary cell cultures using hemocytes and phagocytic tissue cells of Locusta migratoria: an application for locust immunity studies. In Vitro Cellular & Developmental Biology-Animal, 52, 100–106.

Duressa, T. F., Vanlaer, R. & Huybrechts, R. 2015. Locust cellular defense against infections: sites of pathogen clearance and hemocyte proliferation. Developmental & Comparative Immunology, 48, 244–253.

Dushay, M. S. 2009. Insect hemolymph clotting. Cellular and molecular life sciences, 66, 2643–2650.

Gillespie, J. P., Burnett, C. & Charnley, A. K. 2000. The immune response of the desert locust Schistocerca gregaria during mycosis of the entomopathogenic fungus, Metarhizium anisopliae var acridum. Journal of Insect Physiology, 46, 429–437.

Grigorian, M. & Hartenstein, V. 2013. Hematopoiesis and hematopoietic organs in arthropods. Development genes and evolution, 223, 103–115.

Gupta, A. 1979. Hemocyte types: their structures, synonymies, interrelationships, and taxonomic significance. In: Gupta, A. P. (ed.) Insect hemocytes development, forms, functions, and techniques. United States of America: Cambridge University Press.

Hillyer, J. F. 2016. Insect immunology and hematopoiesis. Developmental & Comparative Immunology, 58, 102–118.

Hillyer, J. F. & Strand, M. R. 2014. Mosquito hemocyte-mediated immune responses. Current opinion in insect science, 3, 14–21.

Jones, J. C. 1962. Current concepts concerning insect hemocytes. American Zoologist, 209–246.

Jones, J. C. 1977. The circulatory system of insects, United State of America, Thomas Springfield.

King, J. G. & Hillyer, J. F. 2013. Spatial and temporal in vivo analysis of circulating and sessile immune cells in mosquitoes: hemocyte mitosis following infection. BMC biology, 11, 55.

Latchininsky, A. V. 2013. Locusts and remote sensing: a review. Journal of Applied Remote Sensing, 7, 075099–075099.

Lavine, M. & Strand, M. 2002. Insect hemocytes and their role in immunity. Insect biochemistry and molecular biology, 32, 1295–1309.

Liu, Z., Wu, M., Deng, Z., Wang, Q., Chen, H., Zhang, J. P., Launois, M. & Zheng, Z. 1999. Action mode, persistence and control value of Fipronil for rice grasshoppers Oxya (Orthoptera: Cantantopidae). Insect Science, 6, 62–70.

Ren, C., Li, X., Wu, Z. & Zhang, X. 2014. Morphology of the hemocytes of Bryodema nigroptera Zheng. Chinese Journal of Applied Entomology, 51, 540–547.

Ribeiro, C. & Breh Lin, M. 2006. Insect haemocytes: what type of cell is that? Journal of insect physiology, 52, 417–429.

Statsoft, I. 2011. STATISTICA (data analysis software system). Tulsa, USA. 10 ed.: Statsoft, Inc.

Strand, M. R. 2008. The insect cellular immune response. Insect science, 15, 1–14.

Tame, A., Ozawa, G., Maruyama, T. & Yoshida, T. 2018. Morphological and functional characterization of hemocytes from two deep-sea vesicomyid clams Phreagena okutanii and Abyssogena phaseoliformis. Fish & shellfish immunology, 74, 281–294.

Tojo, S., Naganuma, F., Arakawa, K. & Yokoo, S. 2000. Involvement of both granular cells and plasmatocytes in phagocytic reactions in the greater wax moth, Galleria mellonella. Journal of Insect Physiology, 46, 1129–1135.

Urbański, A., Adamski, Z. & RosińSKI, G. 2018. Developmental changes in haemocyte morphology in response to Staphylococcus aureus and latex beads in the beetle Tenebrio molitor L. Micron, 104, 8–20.

Wang, Q., Cui, Z., Wang, Y., Sheng, X. & Zhang, X. 2011. Comparison among three staining methods to hemocytes of Oxya chinensis. Chinese Journal of Applied Entomology, 48, 841–844.

Wu, G. & Yi, Y. 2015. Effects of dietary heavy metals on the immune and antioxidant systems of Galleria mellonella larvae. Comparative Biochemistry and Physiology Part C: Toxicology & Pharmacology, 167, 131–139.

Yu, C.-H., Yang, H.-Y., Kim, W.-K. & Kim, C.-W. 1977. An Ultrastructural Study on Larval Hemocytes of Acrida cinerea Thunberg. Applied Microscopy, 7, 13–20.

Yu, Y., Cao, Y., Xia, Y. & Liu, F. 2016. Wright–Giemsa staining to observe phagocytes in Locusta migratoria infected with Metarhizium acridum. Journal of invertebrate pathology, 139, 19–24.

Zachary, D. & Hoffmann, D. 1984. Lysozyme is stored in the granules of certain haemocyte types in Locusta. Journal of insect physiology, 30, 405–411.

Zhang, C. & Huang, Y. 2008. Complete mitochondrial genome of Oxya chinensis (Orthoptera, Acridoidea). Acta Biochimica et Biophysica Sinica, 40, 7–18.

